# Coordinated GlcNAc transport couples nutrient acquisition to invasive growth and virulence *in Candida albicans*

**DOI:** 10.64898/2026.10.15.759338

**Authors:** Lanxin Li, Christopher Adamson, Evan Wei Long Ng, Yuan Qiao

## Abstract

*Candida albicans* is a prevalent gut commensal and opportunistic fungal pathogen whose transition from intestinal colonization to invasive infection is influenced by environmental cues. *N*-acetylglucosamine (GlcNAc), abundant in the gut from host and microbial sources, serves not only as a carbon source but also as a potent signal that promotes hyphal morphogenesis and virulence. However, how *C. albicans* coordinates GlcNAc uptake to regulate these pathogenic responses remains incompletely understood. Here, we identify Hgt1, a canonical glucose transporter, as an alternative GlcNAc transporter that cooperates with the GlcNAc-specific transporter Ngt1 to promote fungal morphogenesis and virulence. Simultaneous deletion of *HGT1* and *NGT1* markedly impairs GlcNAc-induced hyphal morphogenesis and virulence-associated gene expression. Mechanistically, Hgt1 facilitates the initial uptake of GlcNAc, promoting Ngt1 induction and subsequent robust hyphal growth, as evidenced by delayed Ngt1 expression and hyphal formation in cells lacking *HGT1*. These findings reveal a coordinated uptake mechanism involving two differentially regulated sugar transporters that enables *C. albicans* to respond efficiently to GlcNAc availability. Importantly, disruption of GlcNAc uptake reduces fungal dissemination to distal organs and associated host inflammatory responses in a mouse intestinal colonization model under GlcNAc-supplemented conditions. Together, our findings uncover a previously unrecognized mechanism by which *C. albicans* coordinates sugar transport to drive nutrient-induced pathogenicity, highlighting GlcNAc uptake as a potential target for limiting fungal virulence and systemic dissemination.

## Introduction

*Candida albicans*, a common human fungal commensal that colonizes mucosal surfaces such as the gastrointestinal (GI) tract, can cause invasive and life-threatening candidiasis when host immunity or microbial homeostasis is disrupted.^1–3^ Central to this transition is *C. albicans*’ ability to switch between yeast and hyphal forms. Hyphal morphogenesis enables *C. albicans* to breach mucosal barriers, invade host tissues, and express a wide range of virulence-associated factors.^4, 5^ In the GI tract, this morphogenetic plasticity unfolds in a dynamic environment where the fungus must integrate multiple signals, including host-derived molecules, microbial metabolites, physicochemical cues, and nutrient availability.^5–7^ How *C. albicans* combines these inputs to coordinate nutrient acquisition with morphogenesis remains poorly understood.

*C. albicans* encounters a highly dynamic carbon landscape in the GI tract, shaped by both the host and resident microbiota.^8^ Among the available carbon sources, *N*-acetylglucosamine (GlcNAc) is particularly prominent, arising from continuous bacterial cell wall remodeling and host glycoconjugates such as mucin, from which the fungal β-N-acetylglucosaminidase Hex1 can release GlcNAc.^9–11^ Consistent with GlcNAc availability in the gut, gastrointestinal-induced transition (GUT) cells of *C. albicans* exhibit upregulation of genes involved in GlcNAc utilization.^12^ GlcNAc, however, is more than a carbon source; it also serves as a potent environmental signal that promotes hyphal morphogenesis,^13, 14^ linking nutrient availability to a morphogenetic program that contributes to virulence. The ability of *C. albicans* to sense and appropriately respond to GlcNAc may therefore be an important determinant of the balance between commensalism and pathogenicity.^11^

The response to GlcNAc is orchestrated by Ngs1, an intracellular GlcNAc sensor that activates transcriptional programs encompassing GlcNAc transport, catabolism, and hypha-associated genes.^15^ Notably, Ngs1-dependent signaling has distinct effects on the host-associated lifestyles of *C. albicans*, promoting virulence while compromising aspects of commensal fitness.^11^ These findings establish GlcNAc sensing as an important mechanism by which *C. albicans* responds to its host environment, but also suggest that GlcNAc transport may function as an upstream determinant of this response. Because extracellular GlcNAc must enter the cell to be sensed intracellularly by Ngs1, its uptake may govern the timing and effects of GlcNAc-induced signaling. *N*-acetylglucosamine transporter 1 (Ngt1) is a well-characterized GlcNAc transporter and a direct transcriptional target of Ngs1,^15, 16^ but evidence from the *ngt1Δ/Δ* mutant suggests other transporters also contribute to GlcNAc uptake.^15–17^ Identifying these transport pathways is essential for understanding how GlcNAc acquisition is coupled to downstream signaling.

Here, we identify Hgt1, a high-affinity glucose transporter, as an alternative GlcNAc transporter. We show that Hgt1-mediated GlcNAc uptake provides an initial signal that facilitates the subsequent induction of the dedicated GlcNAc transporter Ngt1, establishing a transporter hierarchy that supports GlcNAc acquisition and signaling in a manner that is also modulated by glucose availability. Because Ngs1-dependent GlcNAc signaling favors virulence at some cost to commensal fitness, this cascade may allow *C. albicans* to calibrate its GlcNAc response and morphogenetic commitment according to the surrounding carbon environment. Overall, our findings reveal an unexpected role for a canonical glucose transporter in GlcNAc sensing and suggest that transporter hierarchy is a previously unrecognized layer of nutrient-dependent control of fungal morphogenesis and host adaptation.

## Results

### Reported *ngt1Δ/Δ* phenotypes suggest alternative GlcNAc uptake mechanisms in *Candida albicans*

ΔNgt1, identified by Konopka and colleagues in 2007, is the first eukaryotic high-affinity GlcNAc transporter.^16^ Ngt1 is highly specific for GlcNAc transport and is selectively induced by it. Accordingly, the *ngt1*Δ/Δ mutant is unable to utilize GlcNAc as the sole carbon source and exhibits markedly impaired GlcNAc-induced hyphal morphogenesis, establishing GlcNAc uptake as a prerequisite for GlcNAc-dependent signaling and morphogenesis.^16^

However, several lines of evidence indicate that Ngt1 is not the only route of GlcNAc entry. First, the original study showed that although *ngt1*Δ/Δ cells are defective in hyphal growth at 1 mM GlcNAc but largely recover filamentation at 100 mM.^16^ Given that intracellular GlcNAc is required for hyphal signaling, this concentration-dependent rescue suggests that additional uptake mechanisms can compensate for the loss of Ngt1 when extracellular GlcNAc concentration is high. In addition, GlcNAc stimulation still triggers a low-level, Ngs1-dependent induction of GlcNAc catabolic genes in the *ngt1*Δ/Δ mutant, implying residual GlcNAc uptake.^15^ Moreover, GlcNAc-based chemical probes are incorporated into *C. albicans* cell-wall chitin through an Hxk1-dependent process that is unaffected by Ngt1 deletion,^17^ further pointing to Ngt1-independent uptake. These observations prompted us to search for alternative GlcNAc transporters, since targeting both Ngt1 and its alternatives could more effectively block GlcNAc uptake, thereby suppressing GlcNAc-dependent signaling and invasive growth in *C. albicans*.

### The high-affinity glucose transporter Hgt1 as a candidate alternative GlcNAc transporter

To identify alternative GlcNAc transporters in *C. albicans*, we drew inspiration from the model yeast *Saccharomyces cerevisiae*, which lacks both dedicated GlcNAc transporters and catabolic pathways, yet imports low levels of GlcNAc through the promiscuous activity of high-affinity hexose transporters (HXTs).^18^ We therefore focused on the *C. albicans* high-affinity glucose transporter (HGT) family, the closest orthologues of *S. cerevisiae* HXTs, as a source of candidate GlcNAc transporters (**Figure 1A**).

**Figure 1.**
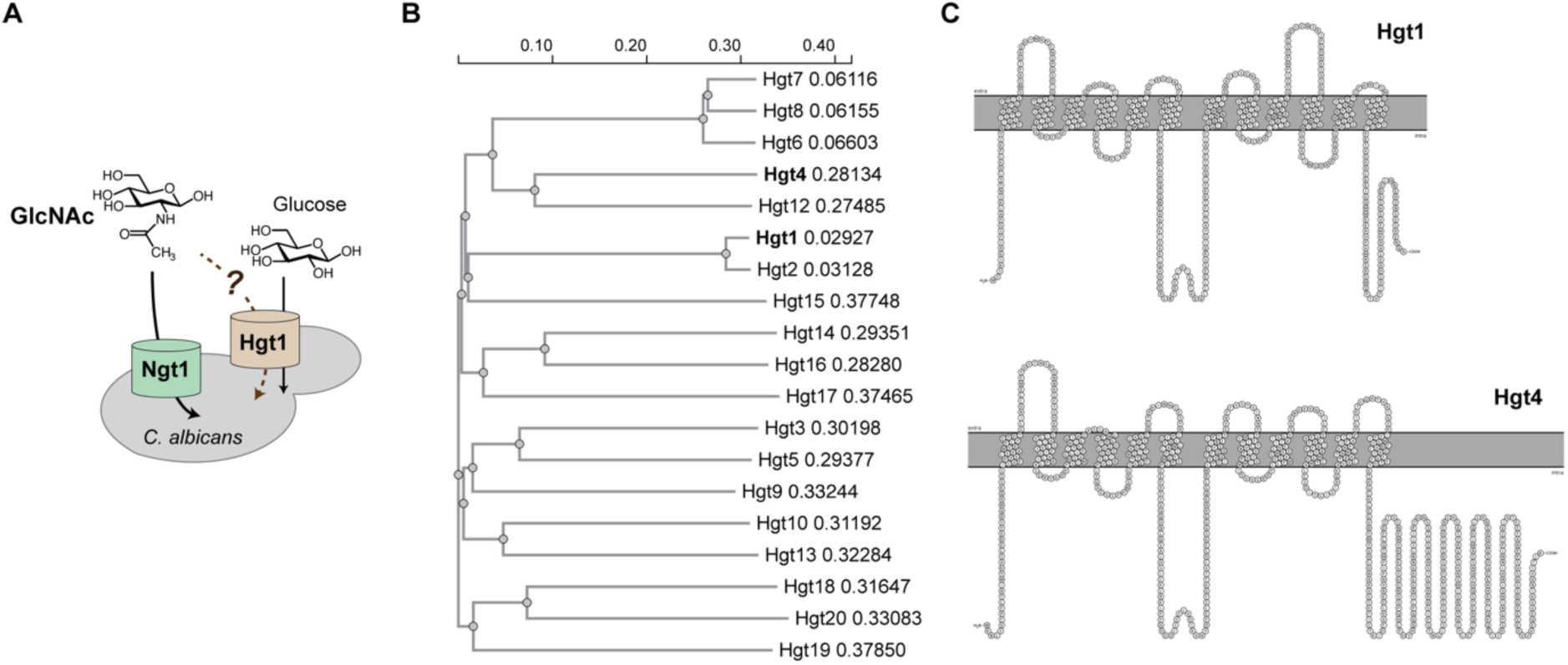
Hgt1 as a candidate alternative GlcNAc transporter. **A.** Schematic illustrating GlcNAc uptake in *C. albicans* mediated by Ngt1, the GlcNAc-specific transporter, and Hgt1, a canonical glucose transporter, which we propose to investigate as an alternative GlcNAc transporter in this study. **B.** Phylogenetic tree of the HGT family in *C. albicans*. **C.** Predicted membrane topology of *C. albicans* Hgt1 and Hgt4 generated using TMHMM and visualized with PROTTER

The *C. albicans* HGT family comprises 20 members of the major facilitator superfamily (MFS), each predicted to contain 12 transmembrane helices (**Figure 1B**).^19^ Among these, Hgt1 and Hgt4 are the best characterized. Hgt1 was the first glucose transporter identified in *C. albicans*, and its heterologous expression in an *S. cerevisiae* strain lacking all HXTs restores glucose uptake.^20^ In contrast, Hgt4 functions as a glucose sensor, which has an extended C-terminal cytoplasmic tail that is highly homologous to those of the *S. cerevisiae* glucose sensors Rgt2 and Snf3 (**Figure 1C**).^21^ Consistent with its regulatory role, the *hgt4Δ/Δ* mutant shows impaired growth on several fermentable sugars, potentially due to the reduced expression of other *HGT* genes encoding functional sugar transporters.^21^ Given that Hgt1 is an established high-affinity glucose transporter and that some glucose transporters in *S. cerevisiae* can also import GlcNAc, we selected Hgt1 as a candidate GlcNAc transporter for further investigation.

### Loss of HGT1 impairs GlcNAc-induced hyphal growth in *ngt1Δ/Δ* under high GlcNAc concentrations

Because the *ngt1Δ/Δ* mutant regains the ability to form hyphae at high GlcNAc concentrations,^16^ we used GlcNAc-induced hyphal growth as an initial readout to test whether Hgt1 deletion impairs this phenotype. To this end, we generated an *hgt1Δ/Δ* single mutant and an *ngt1Δ/Δ hgt1Δ/Δ* double mutant (2KO) for comparison with *ngt1Δ/Δ*. Consistent with previous reports, *ngt1Δ/Δ* showed impaired hyphal growth (<10%) at 2 mM GlcNAc, but this defect was largely rescued at 10 mM, with nearly 80% of cells forming hyphae, comparable to the wildtype cells (**Figure 2A-C**). In contrast, the 2KO mutant exhibited a persistent defect in hyphal formation, with no detectable hyphae at 2 mM and only ∼20% hyphal cells at 10 mM GlcNAc (**Figure 2A–C**). This defect remained evident at 20 mM; however, at 50 mM, hyphal growth was also reduced in wildtype cells under our assay conditions, resulting in no appreciable difference between 2KO and wildtype (**Figure S1**). Importantly, reintroducing a single copy of *HGT1* into 2KO effectively restored its hyphal growth at 10 mM GlcNAc (**Figure 2A-C**), confirming that the hyphal defect resulted from *HGT1* deletion rather than secondary or polar effects. Moreover, the 2KO mutant failed to induce the expression of hypha-specific genes *UME6* and *SAP6* in response to GlcNAc, both of which are closely associated with *C. albicans* virulence *in vivo.*^22, 23^ This defect was also restored by *HGT1* complementation (**Figure 2D-E)**. Taken together, deletion of *HGT1* in the *ngt1Δ/Δ* background (i.e. the 2KO mutant) impairs GlcNAc-induced hyphal growth at high GlcNAc concentrations, consistent with Hgt1 partially compensating for the loss of Ngt1. However, the residual hyphal growth of the 2KO (i.e. ∼20% at 10 mM GlcNAc) suggests that additional transporter(s) contribute to this process.

**Figure 2.**
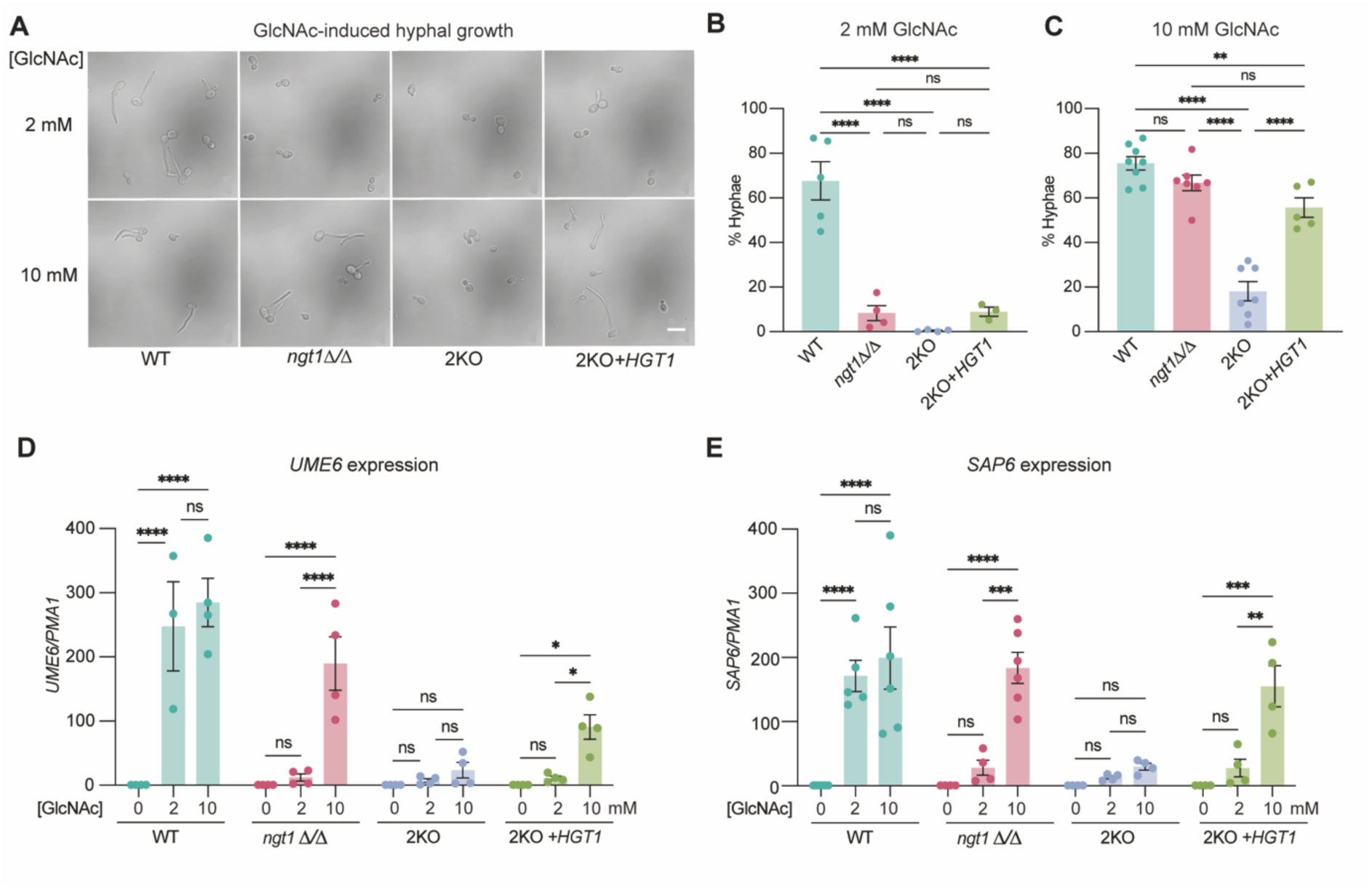
Hgt1 coordinates with Ngt1 to facilitate GlcNAc-induced hyphal morphogenesis in *C. albicans*. **A.** The *ngt1Δ/Δ hgt1Δ/Δ* (2KO) mutant exhibits impaired hyphal growth across a range of GlcNAc concentrations. Representative images showing the morphology of the indicated *C. albicans* strains following incubation with 2 or 10 mM GlcNAc for 4 h. Scale bar, 10 µm. **B-C.** Quantification of the percentage of hyphal cells under the conditions described in **A**. Data represent the mean percentage of hyphal cells ± SEM from ≥3 independent experiments, with ≥150 cells analyzed per experiment. Statistical significance was determined using one-way ANOVA followed by Dunnett’s correction for multiple comparisons. **D-E.** The 2KO mutant exhibits reduced expression of virulence-associated genes in response to GlcNAc. RT-qPCR analysis of the expression of *UME6* (**D**) and *SAP6* (**E**) following incubation with 2 or 10 mM GlcNAc for 4 h in the indicated *C. albicans* mutants. Data represent the mean ± SEM from ≥4 independent experiments. Statistical significance was determined using two-way ANOVA followed by Tukey’s correction for multiple comparisons.

To further define the contribution of Hgt1, we examined the *hgt1*Δ/Δ single mutant. Although its hyphal growth was indistinguishable from the wildtype at 4 h, a time-course analysis revealed a pronounced delay in its hyphal induction after 2 h of GlcNAc exposure. This delay occurred at both 2 mM and 10 mM GlcNAc and was rescued by *HGT1* complementation (**Figure S2A–C**). Thus, even in the presence of the high-affinity transporter Ngt1, loss of Hgt1 delays the hyphal response to GlcNAc. This suggests that Hgt1 may support the initial intracellular accumulation of GlcNAc needed to trigger hyphal initiation, a role distinct from that of Ngt1 in sustained high-affinity uptake. Notably, all mutants showed normal FBS-induced hyphal growth and liquid culture growth curves, indicating GlcNAc-specific hyphal defect rather than a general disruption of hyphal morphogenesis or growth (**Figure S2D–F**).

### Hgt1 mediates uptake of GlcNAc-Ph, a surrogate probe for GlcNAc

To determine whether the hyphal defect in 2KO reflects impaired GlcNAc uptake, we sought to directly assess GlcNAc uptake. However, attempts to quantify intracellular GlcNAc (m/z: 222.10) in *C. albicans* by LC-MS were unsuccessful, possibly due to its rapid catabolism that precludes reliable detection (**Figure 3A-B, S3A**). We therefore turned to a chemical biology strategy using GlcNAc probes as surrogates to monitor GlcNAc uptake. Among a series of synthetic analogues bearing different C1-substiuents, the benzyl derivative GlcNAc-Ph stood out as a suitable probe (**Figure 3C, S3B-D)**. Upon incubation, the intact GlcNAc-Ph (m/z: 312.14), together with a small amount of its phosphorylated form (m/z: 390.10), were readily detected in *C. albicans* lysates using LC-MS (**Figure 3C**). This observation confirms the cellular uptake of GlcNAc-Ph and demonstrates that it severs as a suitable substrate for subsequent phosphorylation by Hxk1, the intracellular GlcNAc kinase that catalyzes the first step of the GlcNAc catabolic pathway. Of note, C1-substituted GlcNAc analogues failed to induce hyphal growth (**Figure S3E**), suggesting disrupted GlcNAc recognition or downstream hyphal signalling. Nevertheless, the robust intracellular accumulation of GlcNAc-Ph supports its use as a surrogate for probing GlcNAc uptake in *C. albicans*.

**Figure 3.**
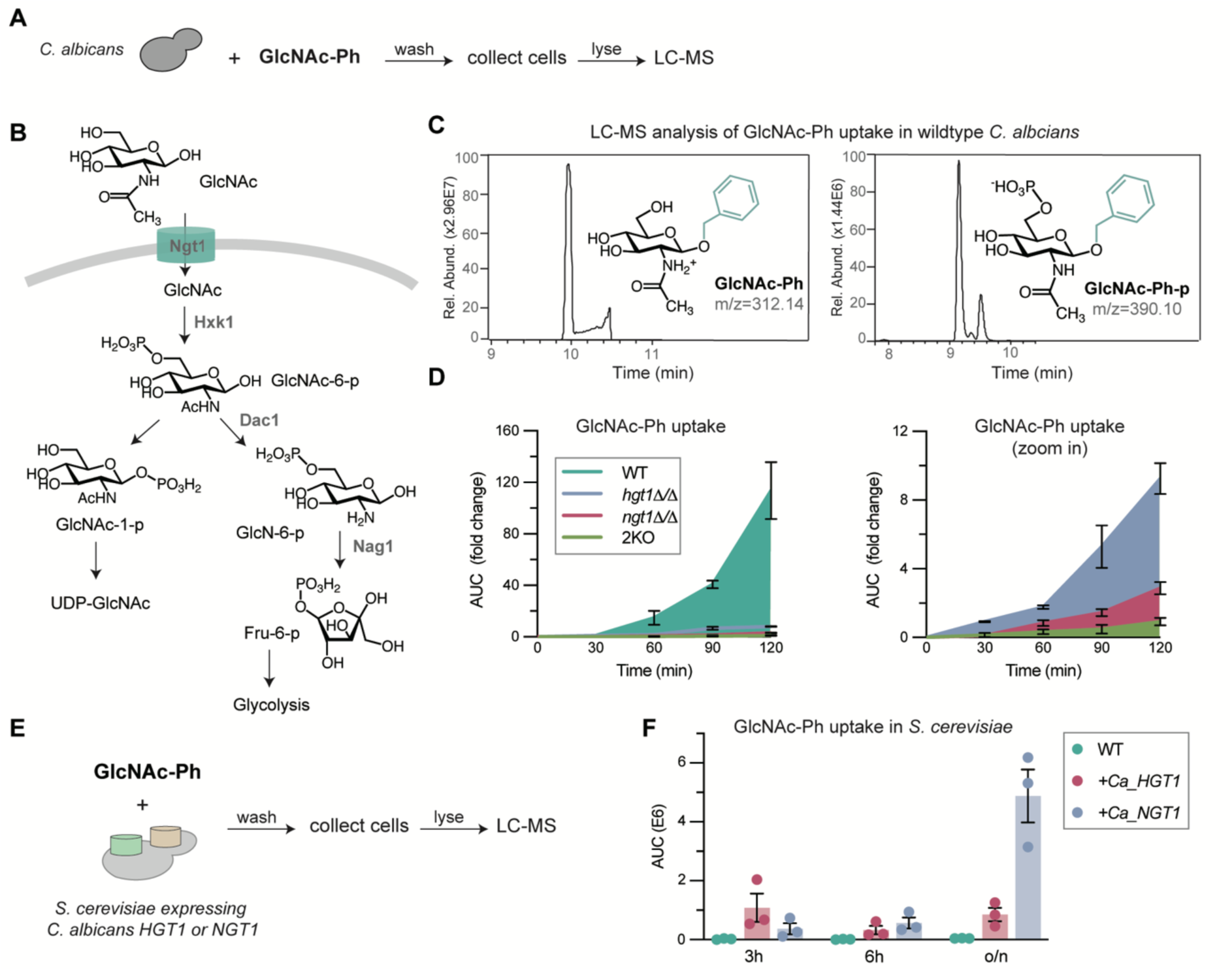
Hgt1 directly contributes to GlcNAc-Ph uptake in coordination with Ngt1. **A.** Schematic illustrating the LC-MS workflow used to assess GlcNAc-Ph probe uptake in *C. albicans*. Briefly, *C. albicans* cells were incubated with 1 mM GlcNAc-Ph for the indicated time points, followed by cell isolation and LC-MS analysis. **B.** Schematic illustrating GlcNAc uptake and catabolism in *C. albicans*. Following internalization by Ngt1, GlcNAc is sequentially phosphorylated by Hxk1 to generate GlcNAc-6-phosphate, deacetylated by Dac1 to generate glucosamine-6-phosphate, and deaminated by Nag1 to generate fructose-6-phosphate, which enters glycolysis for energy production. GlcNAc-6-phosphate can also be converted to GlcNAc-1-phosphate for UDP-GlcNAc biosynthesis. **C.** The GlcNAc-Ph probe is efficiently internalized and phosphorylated by *C. albicans* cells. LC-MS chromatograms of intracellular GlcNAc-Ph and GlcNAc-Ph-phosphate signals detected in lysates of wildtype *C. albicans* cells upon incubation with 1 mM GlcNAc-Ph for 3 h. **D.** Deletion of Hgt1 or Ngt1 severely impairs GlcNAc-Ph uptake. LC-MS quantification of intracellular GlcNAc-Ph in the indicated *C. albicans* strains following incubation with 1 mM GlcNAc-Ph for the indicated time points. The area under curve (AUC) of GlcNAc-Ph LC chromatogram was normalized to that of the 2KO strain at 30 min in each independent experiment. Data represent the mean ± SEM from three independent experiments. **E.** Schematic illustrating the LC-MS workflow used to assess GlcNAc-Ph uptake in *S. cerevisiae* strains heterologously expressing *C. albicans* Hgt1 or Ngt1. **F.** Heterologous expression of *C. albicans* Hgt1 or Ngt1 facilitates GlcNAc uptake in *S. cerevisiae*. LC-MS quantification of GlcNAc-Ph uptake in the indicated *S. cerevisiae* strains following incubation with 1 mM GlcNAc-Ph for the indicated time points. Data represent the mean ± SEM from three independent experiments.

We next compared GlcNAc-Ph uptake in the transporter deletion strains. Both *hgt1Δ/Δ* and *ngt1Δ/Δ* showed a marked reduction in cellular GlcNAc-Ph signals relative to the wildtype, indicating that both transporters contribute to the probe uptake (**Figure 3D**). A detailed comparison revealed a graded pattern, with *hgt1Δ/Δ* exhibiting a slightly higher uptake than *ngt1Δ/Δ* and the 2KO mutant having the lowest level (**Figure 3D**). This finding is consistent with Ngt1 making the predominant contribution, as expected from its established role as a high-affinity GlcNAc transporter, while Hgt1 also contributes substantially to the probe uptake.

The further reduction observed in the 2KO supports their combined roles. Nevertheless, despite their involvement in GlcNAc-Ph uptake, Hgt1 and Ngt1 differed in their contributions to GlcNAc utilization. Unlike *ngt1Δ/Δ*, which was unable to grow with GlcNAc as the sole carbon source, *hgt1Δ/Δ* displayed normal growth on both GlcNAc and glucose (**Figure S4**), likely reflecting the functional redundancy among Hgt-family transporters in sugar acquisition.

To further test whether Hgt1 can directly mediate GlcNAc uptake, we heterologously expressed *C. albicans* Hgt1 and Ngt1 in *S. cerevisiae,* which lacks the GlcNAc catabolic pathway. However, LC-MS detection of exogenous GlcNAc in *S. cerevisiae* was complicated by background metabolites in the cellular matrix, so we again used GlcNAc-Ph as a surrogate probe (**Figure 3E**). After 3 h of incubation, S. *cerevisiae* expressing Hgt1 accumulated readily detectable intracellular GlcNAc-Ph, at levels slightly higher than those in Ngt1-expressing cells. After overnight incubation, however, Ngt1-expressing cells accumulated substantially more GlcNAc-Ph than Hgt1-expressing cells (**Figure 3F**). This time-dependent reversal suggests that the two transporters differ in transport kinetics and capacity, with Hgt1 possibly acting faster early on and Ngt1 sustaining greater accumulation over longer periods, consistent with their proposed complementary roles in *C. albicans*.

### Hgt1 and Ngt1 are differentially regulated in response to GlcNAc

Having established Hgt1 as an alternative transporter that functions alongside Ngt1, we next investigated their transcriptional and translational regulation in *C. albicans*. We constructed Ngt1-GFP and Hgt1-mCherry reporter strains by tagging one chromosomal copy of *NGT1* with GFP and *HGT1* with mCherry, respectively. Following incubation in PBS, Ngt1-GFP fluorescence was undetectable, whereas Hgt1-mCherry showed strong fluorescence at the cell periphery, indicating constitutive Hgt1 expression in the absence of sugar (**Figure 4A**). Supplementation with GlcNAc robustly induced Ngt1-GFP signals, whereas glucose failed to induce it, consistent with the previously reported GlcNAc-specific induction of Ngt1.^16^ In contrast, both glucose and GlcNAc markedly repressed Hgt1-mCherry fluorescence (**Figure 4A**), whereas other sugars metabolites, such as MurNAc and GlcNAc-MurNAc derived from bacterial peptidoglycan had no appreciable effects on Hgt1-mCherry fluorescence (**Figure S5A**).

**Figure 4.**
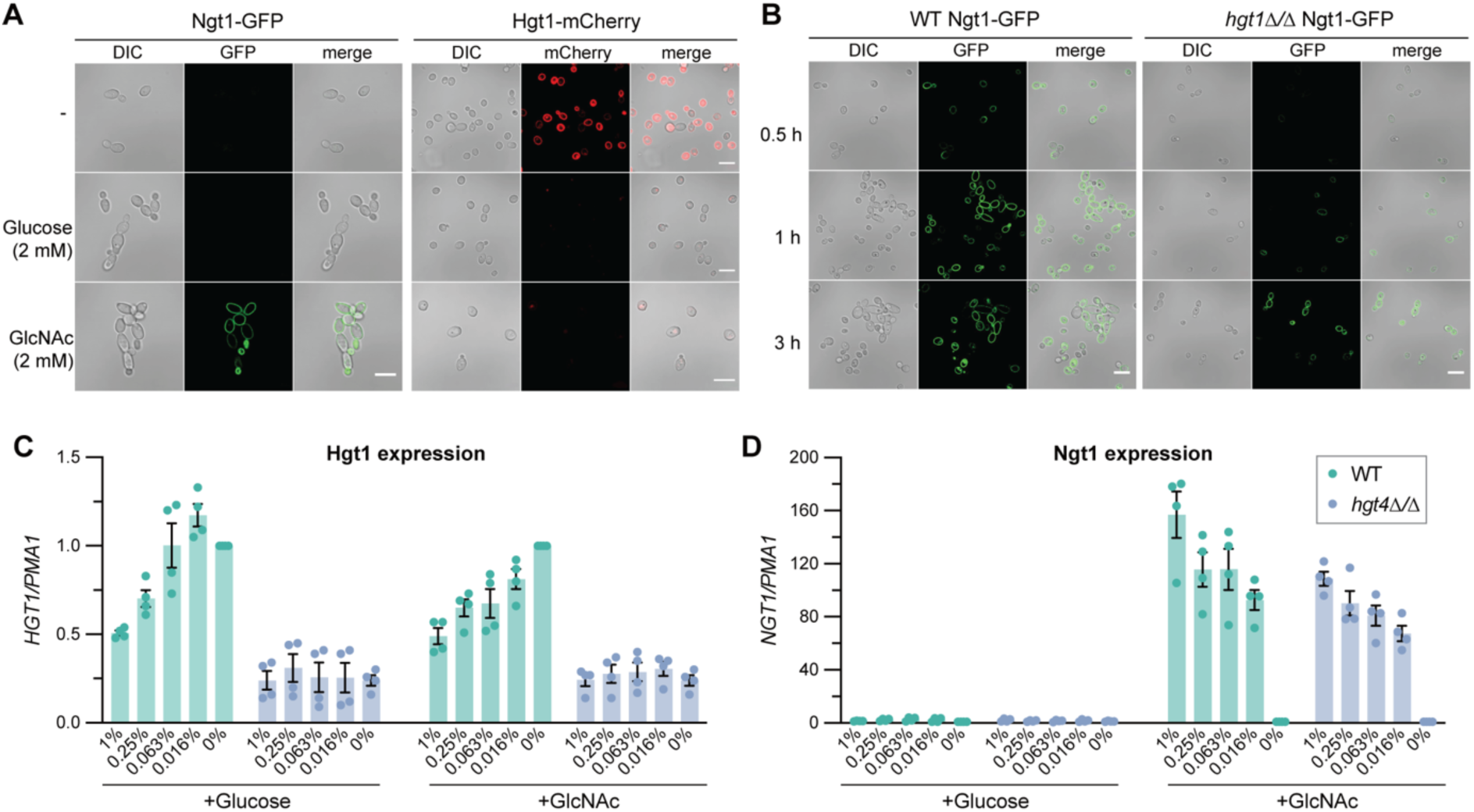
Hgt1 and Ngt1 are differentially regulated by GlcNAc and glucose at the transcriptional and translational levels. **A.** Ngt1-GFP expression is induced by GlcNAc, whereas Hgt1-mCherry expression is suppressed by both GlcNAc and glucose. Representative fluorescence microscopy images showing Ngt1-GFP and Hgt1-mCherry expression and localization in the presence or absence of 2 mM glucose or GlcNAc for 2 h. Scale bar, 10 µm. **B.** Delayed induction of Ngt1-GFP in the *hgt1Δ/Δ* mutant. Representative fluorescence microscopy images showing Ngt1-GFP expression in wildtype and *hgt1Δ/Δ* cells following induction with 0.5 mM GlcNAc for the indicated time points. Scale bar, 10 µm. **C-D.** Both glucose and GlcNAc suppress *HGT1* expression in a dose-dependent manner through Hgt4, whereas GlcNAc induces Ngt1 expression independently of Hgt4. RT-qPCR analysis of *HGT1* (**C**) and *NGT1* (**D**) expression in wildtype and *hgt4Δ/Δ* cells following induction with the indicated concentrations of glucose or GlcNAc for 2 h. Data represent the mean ± SEM from four independent experiments.

The constitutive localization of Hgt1 at the cell surface in the absence of sugar is consistent with its proposed role in facilitating initial GlcNAc uptake, thereby promoting intracellular GlcNAc accumulation and subsequent Ngt1 induction. To test this model, we compared the kinetics of Ngt1-GFP induction in the wildtype and *hgt1Δ/Δ* background in a time-course experiment. Unlike the wildtype strain, in which Ngt1-GFP signal was induced as early as 30 min following GlcNAc exposure, the *hgt1Δ/Δ* mutant showed significantly delayed Ngt1-GFP induction, with weak signals at 1 h and stronger fluorescence emerging only by 3 h. (**Figure 4B**). Notably, the delayed Ngt1 induction in the *hgt1Δ/Δ* mutant coincides with its delayed onset of GlcNAc-induced hyphal growth (**Figure S2A-C)**. Together, our findings support a role for Hgt1 in facilitating initial GlcNAc uptake. Loss of Hgt1 delays uptake-dependent Ngt1 induction, resulting a slower hyphal response to GlcNAc.

At the transcriptional level, GlcNAc modestly repressed *HGT1* in a concentration-dependent manner (**Figure 4C**). Because the glucose sensor Hgt4 regulates multiple sugar transporter genes, including *HGT1*, we next asked whether Hgt4 mediates this repression.^21^ Indeed, deletion of *HGT4* abolished both glucose- and GlcNAc-dependent repression of *HGT1*. Basal *HGT1* expression is also lower in the *hgt4Δ/Δ* mutant, suggesting that Hgt4 supports both basal and sugar-responsive regulation of *HGT1* (**Figure 4C**). Consistent with transcript data, sugar-dependent repression of Hgt1-mCherry at the protein level was likewise lost in the absence of *HGT4* (**Figure S5B**). Furthermore, *hgt4Δ/Δ* phenocopied the delayed GlcNAc-induced hyphal growth of *hgt1Δ/Δ*, further supporting a role for Hgt4 in *HGT1* regulation (**Figure S5C–D**). As a control, *HGT12*, an established Hgt4-regulated target, showed similar Hgt4-dependent transcriptional responses to glucose and, unexpectedly, to GlcNAc (**Figure S5E**). In contrast, GlcNAc-induced *NGT1* expression was unaffected by *HGT4* deletion (**Figure 4D**), as expected because Ngs1 is the dedicated GlcNAc sensor that regulates *NGT1*^15^. Together, these findings indicate that *HGT1* and *NGT1* are regulated through distinct sugar-sensing pathways, which shape GlcNAc-dependent signaling and morphogenesis in *C. albicans*.

### GlcNAc induces endocytosis of both Ngt1 and Hgt1

In yeast, ligand-induced endocytosis of nutrient transporters is a key regulatory mechanism that adapts nutrient uptake to environmental availability.^24, 25^ Accordingly, Ngt1 undergoes endocytosis in response to excess GlcNAc.^26^ We therefore investigated whether Hgt1 is also subject to GlcNAc-induced endocytosis. Upon incubation in sugar-free PBS, Hgt1-mCherry localized predominantly to the plasma membrane but progressively redistributed to the vacuole during prolonged exposure to GlcNAc or glucose, consistent with sugar-induced endocytosis (**Figure S6A-B**).

To directly compare the trafficking of Hgt1 and Ngt1 in response to GlcNAc, we constructed a dual reporter strain co-expressing Ngt1-GFP and Hgt1-mCherry (**Figure 5A**). Neither signal was detectable in overnight YPD cultures. However, upon transfer to sugar-free PBS, Hgt1-mCherry fluorescence emerged at the plasma membrane, whereas Ngt1-GFP was undetectable (**Figure 5B**). Upon GlcNAc addition, Ngt1-GFP was induced at the plasma membrane within 30 min (**Figure 5C**), at which point Hgt1-mCherry was still largely at the cell surface, resulting in clear co-localization of the two transporters (**Figure 5C**). After 3 h of GlcNAc exposure, more than 70% of both Hgt1-mCherry and Ngt1-GFP had relocated from the plasma membrane to the vacuole, indicating GlcNAc-induced endocytosis of both transporters (**Figure 5D**). Quantification showed no significant difference between Hgt1 and Ngt1 in the extent or kinetics of GlcNAc-induced endocytosis (**Figure 5D, F**). In contrast, glucose induced endocytosis of Hgt1-mCherry but did not alter the surface localization of Ngt1-GFP (**Figure 5E**), consistent with their distinct ligand specificities.

**Figure 5.**
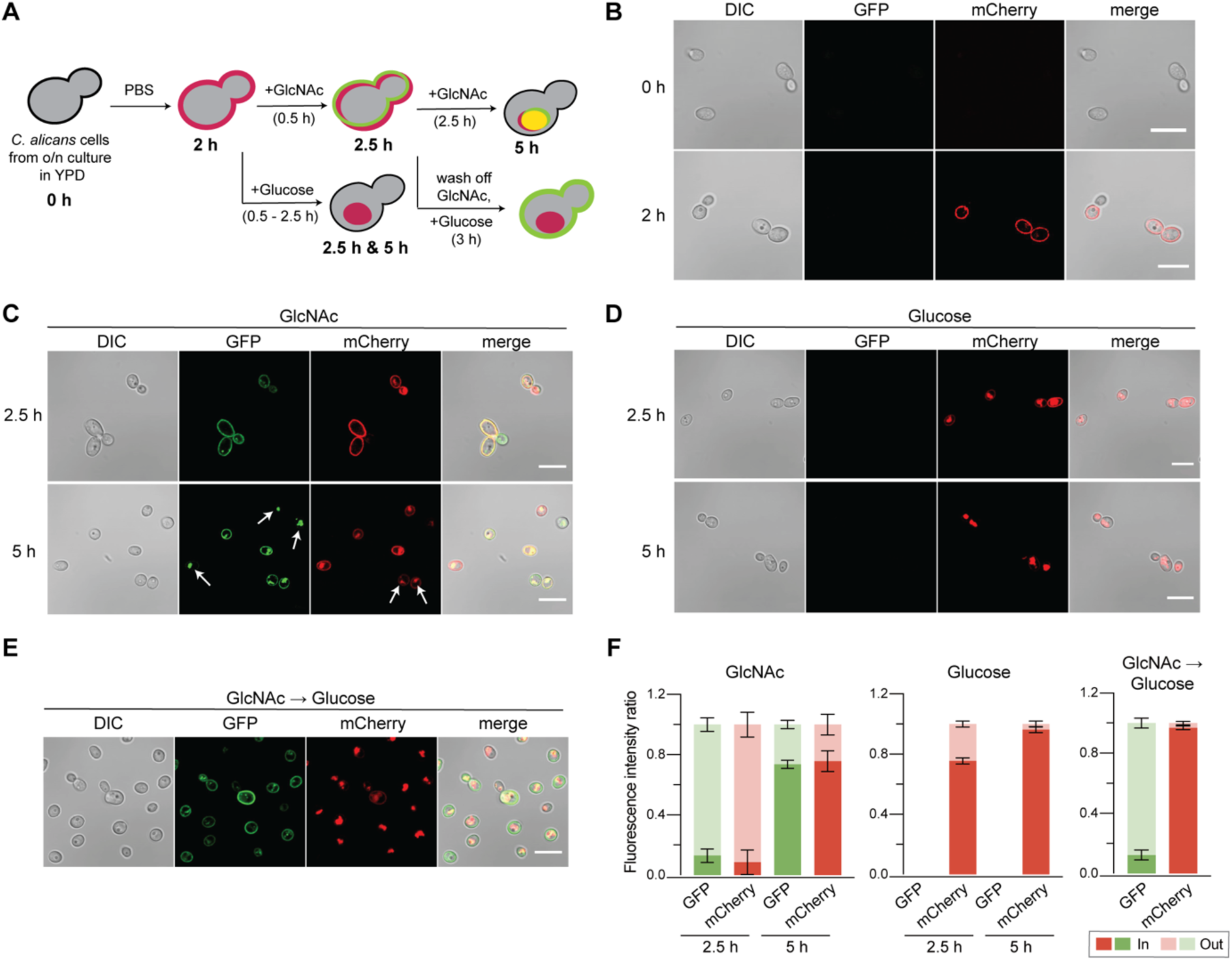
Sugar-induced endocytosis of Hgt1 and Ngt1. **A.** Schematic illustrating the workflow for the ligand-induced endocytosis assay. Briefly, *C. albicans* cells harvested from overnight cultures were washed and incubated in PBS for 2 h to promote Hgt1-mCherry localization at the cell periphery. Cells were then incubated with either 2 mM GlcNAc or glucose for 30 min, followed by continued incubation with the representative sugar for up to 5 h in total. In a separate condition, cells exposed to GlcNAc for 0.5 h were washed to remove GlcNAc and subsequently incubated with 2 mM glucose for an additional 3 h incubation. **B-E.** Representative fluorescence microscopy images showing the expression and localization of Ngt1-GFP and Hgt1-mCherry at 0 and 2 h (**B**), following exposure to GlcNAc (**C**) or glucose (**D**) at 2.5 and 5 h, and following GlcNAc removal and subsequent replacement with glucose (**E**). White arrows indicate endocytosed proteins in *C. albicans* vacuoles. Scale bar, 10 µm. **F.** Quantification of the ratio of intracellular to cell-surface fluorescence signals in the images shown in **B-E,** with the total fluorescence intensity of each cell normalized to 1. Fluorescence intensity was quantified using ImageJ. Data represent the mean fluorescence intensity ratio ± SEM from at least five cells

We next compared the sensitivity of Hgt1 endocytosis to glucose and GlcNAc by quantifying the relative Hgt1-mCherry signals at the vacuole and cell surface. In the dual reporter strain, exposure to 2 mM glucose for 30 min induced greater Hgt1-mCherry internalization than exposure to 2 mM GlcNAc (**Figure 5C–D, F**), suggesting more rapid internalization in response to glucose. Moreover, following 3 h incubation, more than 80% of Hgt1 was internalized at 0.25 mM glucose, whereas a 40-fold higher concentration of GlcNAc (10 mM) was required to achieve comparable internalization (**Figure S6C–E**). Together, these results indicate that Hgt1 endocytosis occurs more rapidly and at lower concentrations in response to glucose than to GlcNAc.

### Hgt4 is required for glucose-mediated suppression of GlcNAc-induced hyphal growth

Although glucose alone does not induce *NGT1* expression, it suppresses its induction by GlcNAc,^16^ indicating that glucose availability modulates the response of *C. albicans* to GlcNAc at the level of transporter induction. We therefore asked whether this glucose-dependent regulation extends to the downstream hyphal response to GlcNAc. Indeed, an equimolar concentration of glucose significantly attenuated GlcNAc-induced hyphal growth but did not affect FBS-induced hyphal development (**Figure 6A-C** and **S7A**), indicating that glucose specifically interferes with the GlcNAc response rather than broadly impairing hyphal induction.

**Figure 6.**
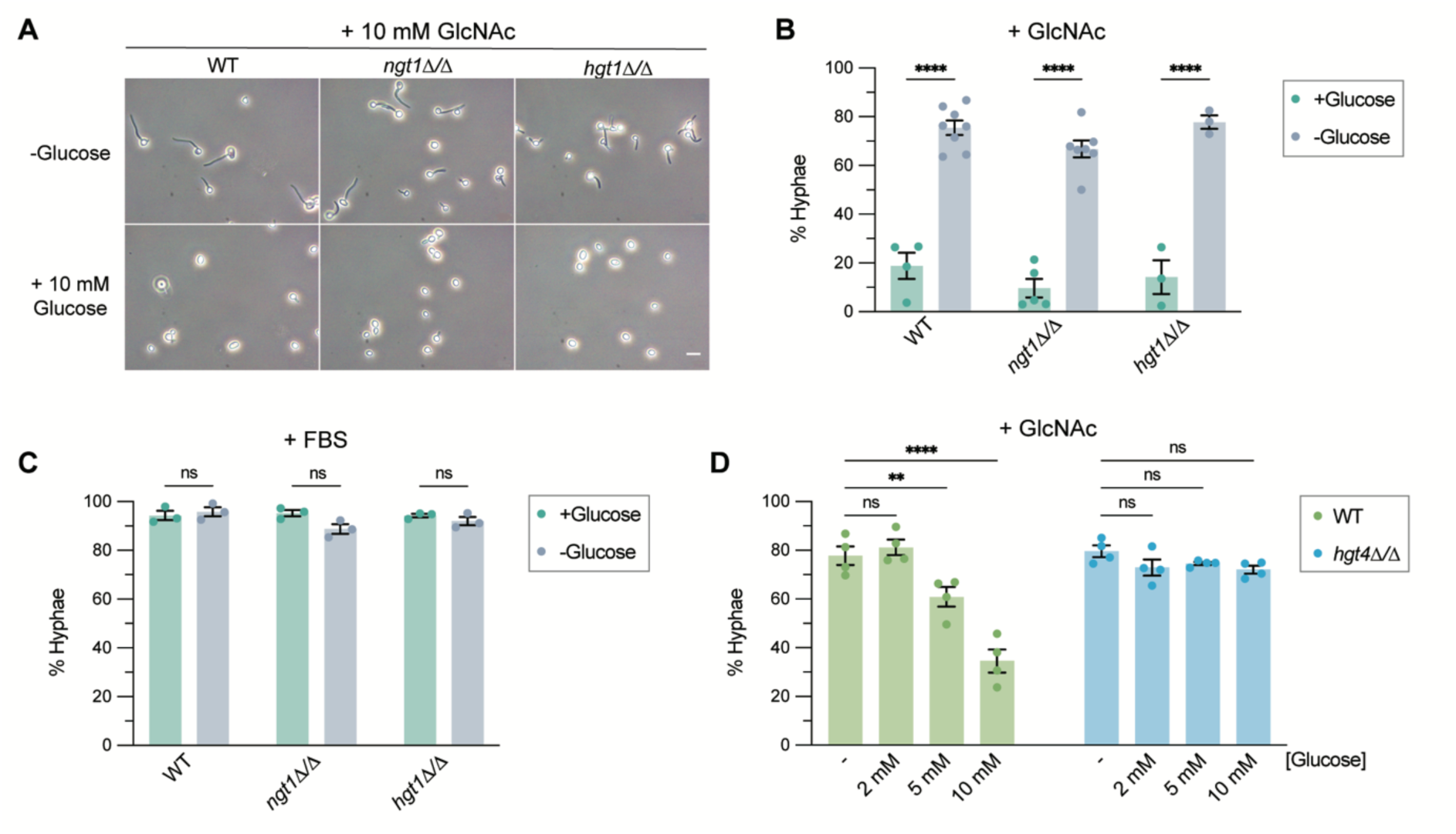
Glucose suppresses GlcNAc-induced hyphal morphogenesis via Hgt4. **A.** Representative microscopy images showing the morphology of the indicated *C. albicans* mutants following induction with 10 mM GlcNAc in the presence or absence of 10 mM glucose. Scale bar, 10 µm. **B.** Quantification of the hyphal cell percentage under the conditions described in **A**. **C.** Glucose does not inhibit FBS-induced hyphal morphogenesis. Quantification of the hyphal cell percentage of wildtype cells following induction with 10% FBS in the presence or absence of 10 mM glucose. **D.** Hgt4 is required for glucose-mediated suppression of GlcNAc-induced hyphal morphogenesis. Quantification of the hyphal cell percentage in wildtype and *hgt4Δ/Δ C. albicans* cells following induction with 10 mM GlcNAc in the presence of the indicated concentrations of glucose. All hyphal quantification data represent the mean percentage of hyphal cells ± SEM from ≥3 independent experiments, with ≥150 cells analyzed per experiment. Statistical significance was calculated using two-way ANOVA followed by Tukey’s correction for multiple comparison

Because glucose impedes Ngt1 induction, we first considered whether it inhibits GlcNAc-induced hyphal growth by reducing Ngt1 expression. However, glucose suppressed GlcNAc-induced hyphal growth similarly in the *ngt1Δ/Δ* mutant (**Figure 6A-B**), suggesting that reduced Ngt1 expression is not the sole mechanism. Because Hgt1 and potentially other Hgt-family transporters can compensate for the loss of Ngt1 to sustain GlcNAc-induced hyphal development, we next asked whether Hgt-familly proteins contribute to the suppression. Deletion of *HGT1* alone did not abolish suppression (**Figure 6A-B**), which may reflect functional redundancy among the Hgt-family members. We therefore examined the *hgt4Δ/Δ* mutant, which exhibits reduced basal expression and impaired sugar-responsive regulation of multiple *HGT* genes.^21^ In contrast to the dose-dependent suppression observed in wildtype cells, glucose failed to suppress GlcNAc-induced hyphal growth in the *hgt4Δ/Δ* mutant (**Figure 6D, S7B**). Together, these results show that Hgt4 is required for glucose-mediated suppression of the GlcNAc response and suggest that glucose and GlcNAc signaling are coordinated in *C. albicans*. Such coordination could allow the fungus to tune its morphogenesis to the relative availability of glucose and GlcNAc, which may be particularly relevant in the gastrointestinal tract.

### GlcNAc promotes tissue dissemination and pathogenicity of gut-colonized *C. albicans*

We next examined how GlcNAc availability affects *C. albicans* colonization and virulence in a mouse model, comparing fungal colonization, morphology, and tissue dissemination in mice given either regular drinking water or water supplemented with 5% GlcNAc (**Figure 6A**). GlcNAc supplementation increased *C. albicans* intestinal colonization approximately 3-fold at 24 h, with about 1.2x10^6^ CFU/mg feces recovered from GlcNAc-treated mice compared with 4x10^5^ CFU/mg from controls (**Figure 6B**). This difference diminished by 72 h with no longer statistical significance, consistent with the notion that sustained GlcNAc signaling compromises the long-term gut fitness of *C. albicans.*^11^ Notably, fecal *C. albicans* recovered from GlcNAc-treated mice contained a markedly higher proportion of hyphal cells (∼70%) than that from control mice (∼40%) (**Figure 6C-D**), indicating that elevated luminal GlcNAc promotes filamentation *in vivo*.

Despite its modest effect on colonization, GlcNAc supplementation markedly enhanced systemic dissemination. At 96 h, fungal burdens were significantly higher in the brain, lungs, liver, spleen, and kidneys of GlcNAc-fed mice (**Figure 6E**), exceeding 10^2^ CFU in the liver and kidney, whereas no colonies were recovered from these organs in control mice. The increased tissue fungal burden was accompanied by elevated expression of the pro-inflammatory cytokines *Tnfα*, *Il6* and *Il12a* in the kidneys and colon of GlcNAc-treated mice (**Figure 6F-G, S8A**), suggesting an enhanced host inflammatory response associated fungal dissemination. Together, these findings indicate that elevated GlcNAc availability promotes hyphal morphogenesis and enhances the invasive potential of gut-colonized *C. albicans*, facilitating its systemic dissemination to distal organs.

### GlcNAc uptake-deficient *C. albicans* exhibits reduced gut colonization and systemic dissemination under GlcNAc-replete conditions

To determine whether GlcNAc uptake contributes to *C. albicans* virulence under GlcNAc-replete conditions, we compared wildtype *albicans* with the GlcNAc uptake-deficient 2KO mutant. Mice were orally gavaged with either strain and maintained on 5% GlcNAc throughout the experiment (**Figure 7A**). Under these conditions, wildtype *C. albicans* reached a fecal burden of approximately 1.5 × 10^6^ CFU/mg, whereas the 2KO mutant reached only ∼5 × 10^5^ CFU/mg (**Figure 7B**), comparable to the wildtype burden in mice without GlcNAc supplementation (**Figure 6B**). Consistently, the 2KO mutant also showed a lower burden in the colonic contents (**Figure 7C**). Moreover, it remained predominantly in the yeast form in the gut, with markedly reduced systemic dissemination and lower induction of pro-inflammatory cytokines in host tissues (**Figure 7D-H, S8B**). Together, these findings indicate that efficient GlcNAc uptake allows *C. albicans* to exploit host luminal GlcNAc, promoting intestinal colonization, hyphal morphogenesis, systemic dissemination, and the associated inflammatory response.

**Figure 7.**
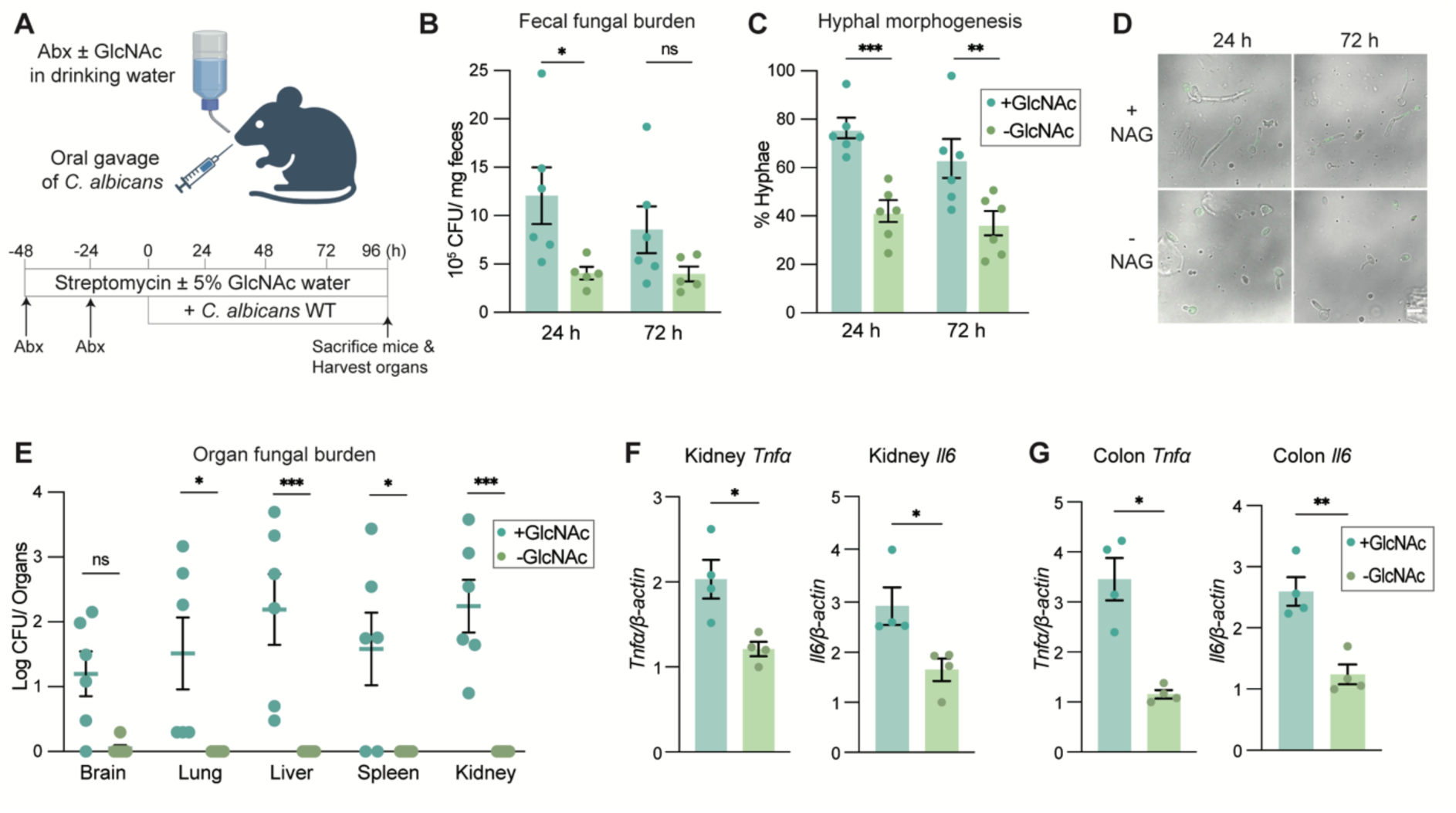
Oral intake of GlcNAc promotes hyphal morphogenesis and tissue dissemination of gut-colonized *C. albicans*. **A.** Schematic overview of the mouse experimental design. Briefly, BALB/c mice (n = 5–6) were orally administered Pen/Strep (penicillin 2,000 units and streptomycin 2 mg/mL) at −48 and −24 h, followed by continuous supplementation of drinking water with 2 mg/mL streptomycin in the presence or absence of 5% GlcNAc throughout the experiment. At 0 h, wildtype *C. albicans* (1 x 10^8^ CFU) was introduced by oral gavage. Mice fecal samples were collected at the indicated time points for analysis of *C. albicans* morphology and fungal burden. Mice were sacrificed at 96 h for organ harvesting and determination of tissue fungal burden. **B.** Oral GlcNAc supplementation slightly increases *C. albicans* gut colonization at the early time point of 24 h, as determined by CFU counts from fecal samples. Data represent the mean ± SEM from n = 6 mice in the +GlcNAc group and n = 5 mice in the −GlcNAc group. Statistical significance was calculated using two-way ANOVA followed by Tukey’s correction for multiple comparisons. **C.** Quantification of the hyphal cell percentage in fecal samples from mice receiving drinking water with or without GlcNAc. Data represent the mean ± SEM from 6 mice per group. Statistical significance was calculated using two-way ANOVA followed by Tukey’s correction for multiple comparisons. **D.** Representative images showing morphology of wildtype *C. albicans* tagged with GFP recovered from mouse feces at the indicated time points. Scale bar, 10 µm. **E.** Quantification of *C. albicans* fungal burden in the indicated organs from mice receiving drinking water with or without GlcNAc. The centre line represents the mean, and error bars indicate ± SEM. Statistical significance was calculated using two-way ANOVA followed by Tukey’s correction for multiple comparisons. **F-G.** Elevated pro-inflammatory cytokine expression in host tissues of *C. albicans*-colonized mice receiving GlcNAc supplementation compared with those without supplementation. RT-qPCR analysis of *Tnfa* and *Il6* in the kidney (**F**) and colon (**G**) of mice receiving drinking water with or without GlcNAc. Data represent the mean ± SEM from 4 mice per group. Statistical significance was calculated using a two-tailed t test.

**Figure 8.**
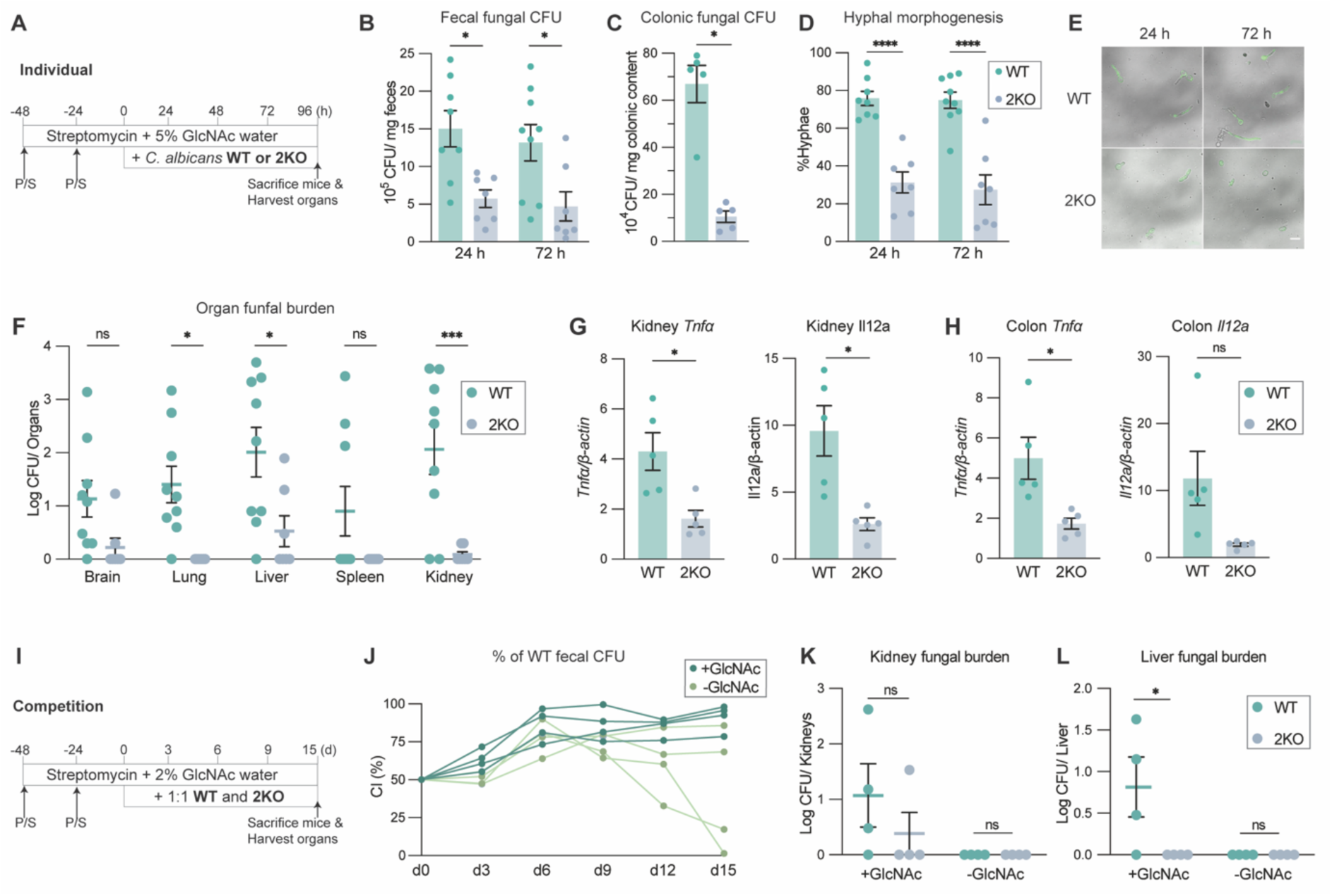
GlcNAc uptake is important for *C. albicans* gut colonization and GlcNAc-associated virulence under GlcNAc-replete conditions. **A.** Schematic overview of the individual colonization mouse model design. Briefly, BALB/c mice (n = 7–8) were orally administered Pen/Strep at −48 and −24 h, followed by continuous supplementation of drinking water with 2 mg/mL streptomycin and 5% GlcNAc throughout the experiment. At 0 h, either wildtype or 2KO *C. albicans* (1 x 10^8^ CFU) was respectively introduced by oral gavage. Fecal samples were collected at the indicated time points for analysis. Mice were sacrificed at 96 h for organ harvesting. **B-C.** Wildtype *C. albicans* exhibits greater gut colonization than the 2KO mutant under GlcNAc-replete conditions, as determined by CFU counts from fecal (**B**) and colonic (**C**) samples. Data represent the mean ± SEM from n = 7–8 mice. Statistical significance was calculated using two-way ANOVA followed by Tukey’s correction for multiple comparisons for fecal fungal CFU (**B**) and a two-tailed t test for colonic fungal CFU (**C**). **D.** Quantification of the percentage of wildtype or 2KO hyphal cells in fecal samples. Data represent the mean ± SEM from 7–8 mice per group. Statistical significance was calculated using two-way ANOVA followed by Tukey’s correction for multiple comparisons. **E.** Representative fluorescence microscopy images showing morphology of GFP-labeled wildtype or 2KO *C. albicans* cells recovered from mouse feces at the indicated time points. Scale bar, 10 µm. **F.** Quantification of wildtype or 2KO *C. albicans* fungal burden in the indicated organs. The center line represents the mean, and error bars indicate ± SEM. Statistical significance was calculated using two-way ANOVA followed by Tukey’s correction for multiple comparisons. **G-J.** Elevated pro-inflammatory cytokine expression in host tissues of mice colonized with wildtype *C. albicans* compared with those colonized with the 2KO mutant. RT-qPCR analysis of *Tnfa* and *Il12a* in the kidney (**G**) and colon (**H**) of mice orally inoculated with either strain. Data represent the mean ± SEM from 5 mice per group. Statistical significance was calculated using a two-tailed t test. **I.** Schematic overview of the competition colonization mouse model design. Briefly, BALB/c mice (n = 4) were orally administered Pen/Strep at −48 and −24 h, followed by continuous supplementation of drinking water with 2 mg/mL streptomycin in the presence or absence of 2% GlcNAc throughout the experiment. At 0 h, wildtype and 2KO *C. albicans* were mixed at a 1:1 ratio (1 x 10^8^ CFU) and introduced by oral gavage. Fecal samples were collected at the indicated time points for analysis. Mice were sacrificed at day 15 for organ harvesting. **J.** Wildtype *C. albicans* outcompetes the 2KO mutant under GlcNAc-replete conditions. The competitive index (CI) was shown as the proportion of the wildtype strain to the total. **K-L.** GlcNAc specifically promotes tissue dissemination of wildtype *C. albicans* but not the 2KO mutant in the competition-colonization model. Quantification of wildtype and 2KO *C. albicans* fungal burden in the kidney (**K**) and colon (**L**) in the presence or absence of GlcNAc. The center line represents the mean, and error bars indicate ± SEM. Statistical significance was calculated using two-way ANOVA followed by Tukey’s correction for multiple comparisons.

To directly compare the fitness of the two strains, we established a competitive colonization model in which wildtype and 2KO were mixed at a 1:1 ratio prior to oral gavage. Mice were then given either regular drinking water or water supplemented with 2% GlcNAc (**Figure 7I**). In the absence of GlcNAc supplemented, the relative abundance of the two strains varied considerably among mice, potentially reflecting individual differences in endogenous luminal GlcNAc availability (**Figure 7J**).^27, 28^ In contrast, under GlcNAc-replete conditions, the wildtype strain consistently outcompeted the 2KO mutant in gut and exhibited greater dissemination to the liver and kidneys (**Figure 7K-L**). These findings highlight the importance of efficient GlcNAc uptake for intestinal fitness and systemic dissemination of *C. albicans*. Collectively, our data demonstrate that elevated luminal GlcNAc preferentially favors *C. albicans* capable of efficient GlcNAc uptake, thereby enhancing intestinal fitness and promoting systemic dissemination *in vivo*.

## Discussion

In this study, we uncovered an unexpected layer of GlcNAc transport regulation in *C. albicans*. The persistence of GlcNAc-induced hyphal morphogenesis in the *ngt1Δ/Δ* cells, which lack the high-affinity GlcNAc transporter Ngt1, suggested additional routes for GlcNAc uptake.^16^ We identified Hgt1, a high-affinity glucose transporter, as one such route, potentially enabled by the structural similarity between glucose and GlcNAc. Consistently, simultaneous deletion of *HGT1* and *NGT1* (2KO) markedly impaired GlcNAc-induced hyphal growth, highlighting the cooperative roles of these two transporters in supporting GlcNAc-mediated filamentation.

Our findings support a model in which Hgt1 mediates the initial uptake of GlcNAc, enabling its intracellular recognition by Ngs1 and subsequent induction of Ngt1 expression.^15^ This sequential process may facilitate a transition form Hgt1-mediated GlcNAc uptake to sustained GlcNAc influx through Ngt1, thereby promoting efficient morphogenesis. Consistent with this mechanism, *hgt1Δ/Δ* cells showed delayed Ngt1 induction and a pronounced delay in GlcNAc-induced hyphal initiation. The eventual recovery of hyphal growth in *hgt1Δ/Δ* cells suggests that alternative uptake mechanism can compensate for the loss of Hgt1 over time. Together, these findings support that Hgt1 functions not merely an alternative GlcNAc transporter, but as an early component of a coordinated transport system that facilitates the timely initiation of GlcNAc-induced hyphal morphogenesis.

The regulation of Hgt1 is also consistent with such a role. Hgt1 was readily detected at the plasma membrane in the absence of sugar, providing access to extracellular GlcNAc before induction of the dedicated transporter Ngt1. However, GlcNAc modestly repressed *HGT1* expression and promoted Hgt1 endocytosis. Notably, Hgt1 internalization in response to GlcNAc was substantially slower and required approximately 40-fold higher GlcNAc concentrations to achieve comparable internalization than glucose. This relatively weak and delayed response to GlcNAc likely allows Hgt1 to remain at the cell surface long enough to provide initial GlcNAc uptake and initiate the GlcNAc-responsive program before Hgt1 is removed from the plasma membrane. Co-opting an existing glucose transporter for initial GlcNAc acquisition may represent an efficient strategy for GlcNAc uptake in *C. albicans*. A related principle, in which basal sugar uptake facilitates the induction of dedicated transport systems, has been described for galactose and maltose in *S. cerevisiae.*^29–31^

Beyond nutrient acquisition, this transport relay may allow *C. albicans* to couple GlcNAc availability with morphogenetic responses. Such coordination could be particularly relevant in the GI tract, where the availability of glucose and GlcNAc varies across anatomical and ecological niches.^32^ For instance, *C. albicans* has been reported to exist predominantly in the yeast form in the mouse small intestine, where glucose is relatively abundant, but exhibits mostly filamentation in the distal gut, where glucose is scarce and GlcNAc is available from microbial and host-derived sources.^9, 12, 23^ These *in vivo* observations align with our *in vitro* findings that, under GlcNAc-rich and glucose-limited conditions, Hgt1-mediated initial GlcNAc uptake facilitates timely Ngt1 induction and promotes hyphal initiation. Conversely, elevated glucose levels suppress GlcNAc-induced morphogenesis. Notably, we found that this inhibitory effect requires Hgt4, a glucose-responsive regulator, suggesting that Hgt4-dependent signalling contributes to the integration of glucose and GlcNAc cues in regulating fungal morphogenesis.

To directly investigate the effects of GlcNAc *in vivo*, we administered GlcNAc to mice. GlcNAc supplementation increased *C. albicans* hyphal formation and enhanced dissemination to distal organs, whereas the GlcNAc uptake-deficient 2KO mutant showed neither response and was outcompeted by wild-type cells under GlcNAc-replete conditions. These results suggest that efficient GlcNAc uptake enables *C. albicans* to take advantage of luminal GlcNAc to promote hyphal morphogenesis and dissemination, potentially at the expense of gut fitness, as previously reported. Interestingly, in the absence of GlcNAc supplementation, the relative competitive fitness of wildtype and 2KO cells varied considerably among individual mice. This variability may reflect differences in GlcNAc availability relative to other carbon sources in the gut,^27, 28^ suggesting that the contribution of GlcNAc uptake to intestinal fitness depends on the local nutrient landscape. More broadly, our results indicate that *C. albicans* responses to GlcNAc are influenced by the availability of other nutrients and that coordinated regulation of sugar transporters may enable the fungus to adjust its morphogenetic response to changes in carbon source availability. Targeting GlcNAc uptake may therefore represent a potential anti-virulence strategy to limit *C. albicans* invasion in GlcNAc-rich environments, particularly those associated with microbiome dysbiosis.

## Materials and Methods

### Ethics statements

Our research complies with all relevant ethical regulations. Animal experiments were approved by the Institutional Animal Care and Use Committee of Nanyang Technological University (Approval No: A24052) and conducted according to ethical and scientific guidelines.

### Growth conditions of Candida albicans and Saccharomyces cerevisiae

*C. albicans* and *S. cerevisiae* strains used in this study are listed in the Supplementary Data 1. *C. albicans* cells were grown at 30 °C, 200 rpm in YPD (2% yeast extract, 1% peptone, 2% glucose) or glucose minimal medium (GMM: 6.7 g/L yeast nitrogen base without amino acids, 5.0 g/L (NH4)2SO4 and 2% glucose) with appropriate supplementation (80 μg/mL uridine, 40 μg/mL arginine, or 40 μg/mL histidine) added when necessary. The solid medium was prepared by adding 2% agar. *S. cerevisiae* cells were grown at 30 °C, 200 rpm in YPD or GMM lacking the appropriate supplementation (800mg/L adenine, 800mg/L leucine,800mg/L uracil, 800mg/L histidine, 1200mg/L lysine, or 800mg/L tryptophan).

### Construct of *C. albicans* gene deletion and complementation mutants

Single deletion mutants (*hgt1Δ/Δ*, *ngt1Δ/Δ*, and *hgt4Δ/Δ*) were constructed as below: using *hgt1Δ/Δ* as an example, ∼500 bp of the *HGT1* promoter region and the last ∼500 bp within the ORF were amplified from SC5314 genomic DNA using primer pairs HGT1_up500 and HGT1_down500 (Supplementary Data 2). The resulting PCR fragments were cloned into the pYGS1033 vector, flanking the URA3 flipper (UFP) cassette located between XhoI and NotI. The resulting knockout cassette was excised with KpnI and SacII and transformed into the *C. albicans* BWP17 strain, generating a heterozygous deletion strain (*hgt1::UFP/HGT1*). To delete the second allele, the UFP marker was replaced by a HIS1 auxotrophic marker cassette and transformed into the *hgt1::UFP/HGT1* strain to generate the homozygous deletion mutant *hgt1::UFP/hgt1::HIS1*. The mutant was complemented with an ARG4 marker using plasmid pYGS1229 (carrying the ARG4 marker), which was linearized with AscI at the GAL4 3′ UTR and integrated into the genome to avoid phenotypic artifacts due to auxotrophic deficiencies. Complete deletion of HGT1 was validated by colony PCR and auxotrophy testing.

To construct the double deletion mutant *hgt1Δ/Δ ngt1Δ/Δ* (2KO), the URA3 marker was first flipped out from the *hgt1::UFP/hgt1::HIS1* strain by culturing in YCB-BSA medium (23.4 g/L yeast carbon base, 4 g/L BSA) for 2 days, followed by selection on GMM medium supplemented with uridine, arginine, histidine, and 0.1% (w/v) 5-fluoroorotic acid (5-FOA), yielding the *hgt1::FRT/hgt1::HIS1* strain. For *NGT1* deletion, ∼500 bp of the promoter region and the last ∼500 bp of the ORF were amplified using primer pairs NGT1_up500 and NGT1_down500 (Supplementary Data 2) and cloned into pYGS1033 as described above. The knockout cassette was excised with KpnI and SacII and transformed into the *hgt1::FRT/hgt1::HIS1* background to generate the heterozygous mutant hgt1::FRT/hgt1::HIS1 *ngt1::UFP/NGT1*. The UFP cassette was then replaced with an ARG4 auxotrophic marker between XhoI and NotI in the NGT1_pYGS1033 plasmid, and transformed into the strain to generate the homozygous double deletion mutant *hgt1::FRT/hgt1::HIS1 ngt1::UFP/ngt1::ARG4*. Complete deletion of *NGT1* was verified by colony PCR and auxotrophy testing.

For complementation of *HGT1*, the ORF along with ∼1000 bp of the native upstream promoter was amplified from SC5314 genomic DNA using primer pair HGT1_comp (Supplementary Data 2) and cloned into the pYGS1460_SAT1 vector between KpnI and XhoI sites. The resulting plasmid, HGT1_p1460, was linearized using AscI and transformed into both *hgt1::UFP/hgt1::HIS1 +ARG4* and *hgt1::FRT/hgt1::HIS1 ngt1::UFP/ngt1::ARG4* strains for complementation.

### Hyphal induction assay

The hyphal induction assay was performed following the previous protocol^1^. Briefly, *C. albicans* overnight cultures grown in GMM with appropriate supplementation were washed twice with sterile dH2O and once with sterile PBS. Cells were then diluted in PBS to a final concentration of 1 x 10⁶ cells mL. Cells were incubated with GlcNAc or GlcNAc derivatives, or with 5% FBS at 37 °C with shaking at 200 rpm for either 2 or 4 hours. To assess the inhibitory effect of glucose, cells were co-incubated with GlcNAc and glucose at the indicated concentrations under the same conditions. Following incubation, cell morphology was assessed at 40x magnification using an Olympus CX43RF ppright Microscope. The percentage of hyphal cells was determined by counting ≥150 cells per sample.

### Gene expression analysis

*C. albicans* overnight cultures grown in GMM with appropriate supplementation were diluted to an OD_600_ of 0.2, washed twice with 1 mL sterile dH2O, once with sterile PBS, and then resuspended in PBS. For quantification of virulence gene expression, cells were incubated with 2 or 10 mM GlcNAc. To assess the expression of sugar transporter genes (*HGT1*, *NGT1*, and *HGT12*) in response to glucose or GlcNAc, cells were incubated with the respective sugars at the indicated concentrations. Cell suspensions were incubated at 37°C with shaking at 200 rpm for 4 h for virulence gene expression analysis and 2 h for sugar transporter gene expression analysis. Following incubation, cells were harvested, washed twice with sterile dH2O, and resuspended in TR buffer (1st BASE). Cell lysis was performed using glass bead beating in a Precellys 24 Touch homogenizer at 6000 rpm for 1 minute, followed by a 1-minute pause on ice, repeated for four cycles. Total RNA was extracted using the PrimeWay Total RNA Extraction Kit (1st BASE), and approximately 500 ng of total RNA was used for cDNA synthesis with the HiScript III RT Supermix (Vazyme). Quantitative real-time PCR (RT-qPCR) was performed using the SsoAdvanced™ Universal SYBR. Green Supermix (Bio-Rad), following the manufacturer’s recommended protocol. Gene-specific primer pairs used in the assay are listed in Supplementary data 3. Relative gene expression levels were calculated using the ΔΔCt method, normalized to *C. albicans PMA1* as the housekeeping gene.

### *C. albicans* growth assays

For liquid growth assay, overnight *C. albicans* cultures grown in YPD were washed twice with PBS and inoculated into fresh YPD (2% yeast extract, 1% peptone, and 2% glucose) or YPN (2% yeast extract, 1% peptone, and 2% GlcNAc) at an initial OD_600_ of 0.1. Culture growth was monitored hourly by measuring the OD_600_ of 1 mL aliquots using a NanoDrop OneC. For solid growth assay, overnight *C. albicans* cultures grown in YPD were washed twice with PBS and adjusted to 1 x 10^6^ cells/mL. Tenfold serial dilutions (1×, 10×, 100×, and 1,000×) were prepared, and 5 µL of each dilution was spotted onto agar plates containing glucose or GlcNAc as the sole carbon source (1% or 0.05%), yeast nitrogen base without amino acids (6.7 g/L), and agar (2%).

### Heterologous expression of *C. albicans* HGT1 or NGT1 in *S. cerevisiae*

*C. albicans HGT1* or *NGT1* ORF was amplified from SC5314 gDNA using primer pairs HGT1_Scomp and NGT1_Scomp (Supplementary Data 2). The resulting fragments were digested with BamHI and XhoI, and subsequently inserted into the p415GPD plasmid. The resulting HGT1_p415GPD or NGT1_p415GPD plasmid was transformed into *S. cerevisiae* DDY904 using the Frozen-EZ Yeast Transformation II™ Kit (Zymo Research). The introduction of CaHGT1 or CaNGT1 was confirmed by colony PCR and auxotroph test.

### GlcNAc uptake assay

*C. albicans* or *S. cerevisiae* overnight cultures grown in YPD were diluted to an optical OD_600_ = 1.0, then washed twice with sterile dH2O and once with HBSS. Cells were resuspended in 10 mL HBSS and incubated with 10 mM native GlcNAc or 1 mM GlcNAc-Ph at 37 °C with shaking at 200 rpm. At designated time points, 2 mL aliquots of the cell suspension were collected, washed twice with ice-cold sterile dH2O, and resuspended in 1 M sorbitol buffer (50 mM Tris-HCl, pH 7.5, 1 M sorbitol, 0.8 M KCl, 10 mM MgSO_4_). To digest the cell wall, Zymolyase-100T (Nacalai Tesque) was added to a final concentration of 50 μg/mL, and the mixture was incubated at 37 °C with shaking at 200 rpm for 30 minutes. The resulting *C. albicans* protoplasts were pelleted by centrifugation at 300 g for 3 minutes, resuspended in 200 µL sterile dH2O, and boiled at 100 °C for 10 minutes to lyse the protoplasts. The lysates were clarified by centrifugation at 16,000 g for 10 minutes, and 30 μL of the supernatant was subjected to high-resolution LC-MS/MS analysis.

LC-MS/MS was performed using a Vanquish Core HPLC system coupled to an Orbitrap Exploris 120 mass spectrometer. Separation was carried out on a Synergi Hydro-RP column (4.6 x 150 mm, 4 μm, 80 Å; Phenomenex). An elution gradient of 5% to 95% acetonitrile (0.04% FA) in water (0.04% FA) was applied over 16 minutes. Detection was carried out in Selected Ion Monitoring (SIM) mode.

### Construct of fluorescent protein labeled Hgt1 and Ngt1 mutants

To generate the HGT1-mCherry fusion construct, the final 1000 bp of the *HGT1* ORF (excluding the stop codon) was amplified by PCR from SC5314 genomic DNA using the HGT1-mCh primer pair (Supplementary data 2). The PCR product was cloned into the pYGS873 vector between the KpnI and XhoI restriction sites. The resulting plasmid, HGT1_p873, was linearized with SnaBI and transformed into the *C. albicans* BWP17 strain using the Frozen-EZ Yeast Transformation II™ Kit (Zymo Research), yielding the strain *HGT1/HGT1::HGT1-mCherry-URA3*; or into the *hgt4Δ/Δ* mutant background to obtain the *hgt4::FRT/hgt4::HIS1 HGT1/HGT1::HGT1-mCherry-URA3* strain.

For the NGT1-GFP construct, the yeast GFP fragment was first amplified by PCR using p873-GFP primer pair (Supplementary data 2) and cloned into the pYGS873 vector between the XhoI and PmeI sites to replace the mCherry sequence, generating the pYGS873_GFP vector. The final 1,000 bp of *NGT1* was incorporated into the pYGS873_GFP plasmid using the same strategy as for *HGT1*, generating the NGT1_pYGS873_GFP plasmid. To facilitate chromosomal integration at the *NGT1* locus, an AscI restriction site was introduced into the 1000 bp *NGT1* sequence by site-directed mutagenesis using the NGT1_Mut primer pair (Supplementary data 2). The resulting plasmid, NGT1_Mut_pYGS873_GFP, was linearized with AscI and transformed into BWP17 to generate the *NGT1/NGT1::NGT1-GFP-URA3* strain, or into the *hgt1Δ/Δ* mutant background to obtain the *hgt1::FRT/hgt1::HIS1 NGT1/NGT1::NGT1-GFP-URA3* strain.

To enable dual fluorescence labeling, the URA3 marker in NGT1_Mut_pYGS873_GFP plasmid was replaced with an ARG4 auxotrophic marker between the MluI and NotI sites, resulting in the NGT1_Mut_pYGS873_GFP_ARG4 plasmid. This plasmid was linearized with AscI and transformed into the *HGT1/HGT1::HGT1-mCherry-URA3* strain to generate the dual-labeled *HGT1/HGT1::HGT1-mCherry-URA3 NGT1/NGT1::NGT1-GFP-ARG4* strain. Successful integration of the fluorescent protein tagged constructs was confirmed by colony PCR, auxotrophic marker selection, and fluorescence microscopy.

### Fluorescence microscopy

Log-phase *C. albicans* cells grown in GMM with appropriate supplementation were washed twice with sterile dH2O and once with sterile PBS, then resuspended in PBS to a final concentration of 3 x 10⁶ cells/mL. For analysis of Hgt1-mCherry and Ngt1-GFP expression, glucose or GlcNAc was added at various concentrations to 50 μL of the cell suspension. Cells were incubated at 30 °C with shaking at 200 rpm for 2 hours. After incubation, cells were pelleted by centrifugation, washed twice with sterile PBS, and resuspended in 10 μL PBS for imaging.

For the endocytosis assay, Hgt1-mCherry expression was first induced by incubating the cells in PBS at 30 °C, 200 rpm for 2 hours. Subsequently, cells were n incubated with either 2 mM GlcNAc or glucose for 30 min, followed by continued incubation with the same sugar for an additional 3 h or removal of GlcNAc and addition of 2 mM glucose for an additional 3 h. Fluorescence imaging was performed using a ZEISS LSM800 confocal microscope with ZEN Blue software. mCherry was visualized using 561 nm excitation and 610/30 nm emission filters, while GFP was visualized using 488 nm excitation and 510/30 nm emission filters. All images were captured using a 63 x oil immersion objective lens and processed with ImageJ.

### Mouse experiment

Female BALB/c mice aged 7–8 weeks (InVivos, Singapore) were housed under non-SPF conditions. The animal rooms were maintained under a 12 h light/12 h dark cycle at 20-22°C and 50-60% relative humidity, with HEPA-filtered air supplied at 15-20 ACPH using 0.3-μm filtration.

To clear the intestinal microbiota before *C. albicans* administration, mice received 0.3 mL of Pen/Strep (2,000 units penicillin and 2 mg/mL streptomycin) solution by oral gavage at −48 and −24 h. For the individual colonization model investigating the effect of GlcNAc, drinking water supplemented with 2 mg/mL streptomycin, with or without 5% GlcNAc, was provided throughout the experiment. At 0 h, mice were orally gavaged with 1 x 10^8^ GFP-labelled wild-type *C. albicans* cells. For the individual colonization model comparing wildtype *C. albicans* with the 2KO mutant, drinking water containing 2 mg/mL streptomycin and 5% GlcNAc was provided throughout the experiment. At 0 h, mice were orally gavaged with 1 x 10^8^ GFP-labelled wildtype or 2KO *C. albicans* cells. For the competitive colonization model, drinking water supplemented with 2 mg/mL streptomycin, with or without 2% GlcNAc, was provided throughout the experiment. At 0 h, mice were orally gavaged with 1 x 10^8^ *C. albicans* cells comprising a 1:1 mixture of GFP-labelled wildtype and non-GFP-labelled 2KO cells.

Fecal samples were collected at the indicated time points to assess fungal burden and cell morphology. For morphological analysis, fecal samples were soaked in sterile PBS for 30 min and mixed using an inoculation loop. A 1 μL aliquot was collected from the white layer between the precipitate and supernatant and examined using a ZEISS LSM 800 confocal microscope with a 63× oil-immersion objective. For fungal burden determination, fecal samples were weighed, resuspended in sterile distilled water, and homogenized with metal beads using a Precellys 24 Touch homogenizer at 6,000 rpm for 1 min. The homogenates were serially diluted and plated onto YPD agar supplemented with Pen/Strep. *C. albicans* CFUs were counted after incubation at 30°C for 48 h. For the competitive colonization model, plates were imaged using the Cy2 fluorescence channel of an ImageQuant 800 system to distinguish GFP-labelled wildtype colonies from non-GFP-labelled 2KO colonies. GFP-labelled wildtype colonies appeared black, whereas non-GFP-labelled 2KO colonies appeared white, allowing the two populations to be enumerated separately.

Mice in the individual colonization models were sacrificed at 96 h after *C. albicans* administration, whereas mice in the competitive colonization model were sacrificed on day 15. The brain, kidneys, lungs, liver and ∼ 1 cm colon were harvested. For fungal burden determination, each organ was immersed in sterile PBS and homogenized with glass beads at 6,000 rpm for 1 min for one or two rounds, as required. The homogenates were plated onto YPD agar, and CFUs were counted after incubation at 30°C for 48 h. For cytokine expression analysis, kidney or colon tissues were immersed in TRIzol reagent (Ambion, Invitrogen) and homogenized with glass beads at 8,000 rpm for 1 min over three to four cycles. The homogenates were stored at -80°C until further processing. Chloroform was added to the tissue-TRIzol homogenates, which were vortexed and incubated for 5 min before centrifugation at 16,000 × g for 10 min at 4°C. The upper aqueous phase containing RNA was collected for subsequent RNA extraction and RT-qPCR analysis, as described in the Gene expression analysis section. Relative gene expression levels were calculated using the ΔΔCt method and normalized to *Mus musculus* β-actin as the housekeeping gene. Gene-specific primer pairs used in the assay are listed in Supplementary data 3.

## Acknowledgments

We thank members of the Qiao lab for the valuable discussions and feedback throughout this project.

## Author contributions

Conceptualization: L.L., Y,Q.

Methodology: L.L., C. A.

Investigation: L.L., C. A., E.W.L.N.

Supervision: Y.Q.

Writing—original draft: L.L., Y,Q.

Writing—review & editing: L.L., C. A., E.W.L.N., Y.Q.

## Data, code, and materials availability

All data needed to evaluate and reproduce the results in the paper are present in the paper and/or the Supplementary Materials. Reagents and materials generated in this study are available upon request to the corresponding author.

